# Serine palmitoyltransferase-mediated de novo sphingolipid biosynthesis is required for normal insulin production and glucose tolerance

**DOI:** 10.1101/2025.05.14.653935

**Authors:** Chloé Amouyal, Shiou Ping Chen, Jessica Denom, Ilyès Raho, Nadim Kassis, Marc Diedisheim, François Mifsud, Justine Lallement, Eleni Georgiadou, Mark Ibberson, Mikael Croyal, Xian C Jiang, Céline Cruciani-Guglielmacci, Hervé Le Stunff, Guy A. Rutter, Christophe Magnan

**Affiliations:** Department of Diabetology, Pitié-Salpêtrière Hospital, AP-HP, Paris; Sorbonne Université, INSERM, NutriOmic Team, Paris, France; Université Paris Cité, CNRS, Unité de Biologie Fonctionnelle et Adaptative, F-75013 Paris, France; Institut Necker-Enfants Malades, INSERM UMR-S1151, Université Paris Cité, 75015 Paris, France; Clinique Saint Gatien Alliance (NCT+), Saint-Cyr-sur-Loire, France; Department of Diabetology, Cochin Hospital, AP-HP, Paris; Section of Cell Biology and Functional Genomics, Department of Medicine, Endocrinology and Metabolism, Imperial College London, London, UK; Vital-IT group, SIB Swiss Institute of Bioinformatics, Amphipole building, Quartier Sorge, University of Lausanne, 1015 Lausanne, Switzerland; Université de Nantes, CHU Nantes, CNRS, INSERM, l’institut du thorax, F-44000 Nantes, France; Department of Cell Biology, SUNY Downstate Health Sciences University, Brooklyn, New York 11203, USA; Institut des Neurosciences Paris-Saclay, CNRS UMR 9197, Université Paris Saclay, France; Lee Kong Chian School of Medicine, Nanyang Technological University, Singapore, Singapore; CR-CHUM, Université de Montréal, Montréal, QC, Canada; Research Institute of McGill University Health Centre, Montréal, QC, Canada

**Keywords:** Beta cell mass, diabetes, ceramides, sphingomyelin

## Abstract

**Aims/Hypothesis:** The importance for normal insulin secretion of ceramide synthesis is unclear. *De novo* ceramide synthesis requires serine palmitoyl transferase, SPT2, encoded by *Sptl2*.

**Methods:** We generated β-cell-selective *Sptl2* null mice by crossing animals with *floxed* alleles to mice expressing *Cre* recombinase from the *Ins1* locus. Metabolic phenotyping, transcriptomic, functional analyses and histology were performed using standard approaches.

**Results:** Islets from *Sptlc2*^ΔIns1^ mice displayed marked alterations in ceramide and sphingomyelin levels: ceramide content: p=0.016 and p=0.109; sphingomyelin content: p=0.016 and p=0.004 in *Sptlc2*^ΔIns1^ vs *Sptlc2*^CTL^ mice under regular and high fat diet, respectively, despite compensatory increases in the expression of enzymes in the salvage and sphingomyelinase pathways. Correspondingly, profound abnormalities were observed in glucose-regulated insulin secretion and glucose tolerance *in vivo*, both on a regular chow and high fat diet. These changes were associated with a drastic (∼80%) lowering in β-cell numbers, and a more minor increase in delta cell numbers. They were also preserved in animals maintained on a ketogenic diet, consistent with a cell autonomous effect on the β-cell. Despite normal glucose-regulated intracellular calcium dynamics and insulin secretion, marked transcriptomic changes were observed in *Sptlc2*^ΔIns1^ mouse islets, with affected GO terms including lysosome organisation and regulation of autophagy. Consistent with roles for compromised SPT2 function in diseased β-cells, *Sptl2* expression in Balbc and DBA2J mouse islets was lowered by a high fat-diet. Moreover, *SPTLC2* mRNA tended to be lower, and *SPTLC1* mRNA was significantly decreased, in islets from human subjects with type 2 diabetes *versus* normoglycemic individuals.

**Conclusions:** Preserved *de novo* ceramide synthesis is required to maintain normal β-cell mass and thus insulin secretion in mice. Therapeutic approaches which seek to target this process systemically using pharmacological SPT2 inhibitors should thus be treated with caution.

**Research in context:** *- What is already known about this subject?:* Ceramides are key components of sphingolipid metabolism. Excess ceramide levels contribute to lipotoxicity and β-cell apoptosis. α-cell-restricted deletion of *Cers2*, which is responsible for the synthesis of very long ceramide chains, alters the insulin content of pancreatic islets and modifies glucose tolerance. Deletion of *Cers 5* or *6*, responsible for the synthesis of the long chains, has no effect.

*- What is the key question?:* What is the importance of *de novo* ceramide synthesis in β-cells for the normal regulation of insulin production and glucose homeostasis?

*- What are the new findings?:* Inhibition of the *de novo* ceramide synthesis pathway in β-cells, achieved by selective deletion of *Sptlc2*, encoding subunit 2 of the serine palmitoyltransferase (SPT) enzyme, induces a major alteration of glucose tolerance and insulin secretion. This is accompanied by a drastic reduction in β-cell mass and islet insulin content. The remaining islets of *Sptlc2*^ΔIns1^ display normal glucose-regulated intracellular calcium dynamics and insulin secretion despite imbalances in ceramide and sphingomyelin levels and substantial transcriptomic changes. Expression of *SPTLC1*, which encodes the other subunit of the SPT heterodimer, is reduced in islets from humans with type 2 diabetes, and a trend is observed towards lowered *SPTLC2* expression. Taken together, these findings highlight the importance of *de novo* ceramide synthesis for normal β-cell survival and function

*- How might this impact on clinical practice in the foreseeable future?:* By suppressing insulin production, global blockade or inhibition of SPT2, achieved with pharmacological approaches which seek to rescue insulin sensitivity in T2D, may be deleterious for glucose tolerance.

## INTRODUCTION

β-cell destruction underlies the pathology of type 1 diabetes (T1D) and may also play a role in type 2 diabetes (T2D) [1]. Changes in the functional identity of the remaining β-cells contribute to both diseases [2, 3].

An increasing number of studies have assigned a particularly deleterious role to ceramide accumulation in the development of insulin resistance and the deterioration of pancreatic β-cell function [4]. Ceramides belong to the sphingolipid family, and are formed by combining a saturated fatty acid, namely palmitate, with sphingosine via an amide bond. De novo synthesis of ceramides takes place in the endoplasmic reticulum (ER). The rate-limiting enzyme is serine palmitoyl transferase (SPT), which is a heterodimer made up of two subunits, including the structural subunit SPT1 (encoded by *Sptlc1*) and the catalytic subunit, which is either SPTLC2 or SPTLC3 (encoded by *Sptlc2 and Sptlc3, respectively)*. Ceramide can also be produced by two other pathways [5]. The degradation pathway of sphingomyelins by sphingomyelinases takes place in different sub-compartments, including the plasma membrane, mitochondria, lysosomes and Golgi apparatus [5]. Finally, the salvage pathway generates ceramides from complex sphingolipids in lysosomes or endosomes [6].

There is a diversity of ceramide species, reflecting the existence of six isoforms of ceramide synthase located in the ER. These are classified according to the length of the fatty acid chain linked to sphingosine [4]. Ceramide species, and particularly the long chain C18:0 and C16:0-ceramides, have been proposed to be important mediators of lipotoxicity [7]. In obesity, ceramide species can accumulate in insulin-sensitive tissues and in β-cells [8]. Ceramide levels increase in the plasma and skeletal muscle of obese mice [9]. In humans, ceramides are also elevated in the liver and blood of T2D patients [9]. In addition, a relationship between circulating ceramide levels and insulin resistance exists in vivo [10] and has been proposed as a predictive biomarker or T2D development [4].

The decrease in pancreatic β-cell mass and the loss of insulin secretion are correlated with an increase in intracellular ceramide levels and SPT expression within the islet of obese diabetic Zucker rats [11]. In pancreatic β-cells, an excess of ceramides, especially long-chain ceramides, appears to induce apoptosis by blocking proliferation or by provoking mitochondrial damage and the subsequent production of reactive oxygen species [12]. The molecular targets of ceramides include downstream mediators of insulin receptor signaling such as IRS1 and Akt2 in responsive tissues [13]. More recently, protein targets of ceramide in insulin resistance have been described such as mitochondrial fission factor, whose activation induced mitochondrial fragmentation [14]. The role of these proteins in ceramide-induced β-cell apoptosis remains to be determined.

In addition to their importance for membrane integrity, ceramides are also important mediators of intercellular signaling and are needed to ensure the correct proportions of other sphingolipid categories [6, 15]. Moreover, it has been proposed that ceramides serve as gauges of free fatty acid excess, favoring their storage as triglycerides and preventing glucose utilization in the liver [16]. Consistent with a beneficial role for ceramides in metabolically-sensitive tissues, selective ceramide species appear to be important in the β-cell for normal glucose homeostasis. Thus, deletion of *CerS2*, either systemically or selectively in the pancreatic β-cell, to limit the production of very long ceramide chains, lowers the insulin content of pancreatic islets and impairs glucose intolerance on both normal and high fat diet [17–20]. In contrast, deletion of *CerS6* or *CerS5* in the β-cell which lowers the synthesis of the long chain forms (C16 or C18-ceramide), has no effect [18]. Interestingly, genome-wide association studies (GWAS) have revealed associations between a single nucleotide polymorphism in *CerS2* and altered glucose homeostasis [21]. CerSs are also known to be central for both *de novo* ceramide synthesis and the salvage pathway [5].

Up to now, the specific role of the initiation of *de novo* ceramide synthesis in β-cells for normal insulin production and the regulation of glucose homeostasis has been unclear. To address this question, we have developed mice deleted selectively in the β-cell for SPT2, and explored the impact on glucose homeostasis.

## METHODS

Detailed methods are presented in Supplemental Data: Extended Methods. All procedures were carried out in accordance with the European Union Directive 2010/63/EU for animal experiments and were approved by the institutional animal care and use committee of the University Paris Cité (CEEA40, authorization <u>#37997-2022022315187624 v7</u>).

### Generation and characterization of pancreatic β-cell-specific *Sptl2* knockout mouse

*Sptlc2*^lox/lox^ mice [22] were crossed with mice in which *Cre* recombinase was knocked in at the *Ins1* locus to confer pancreatic β-cell-specific expression (*Ins1*Cre^+/-^) [23]. Multiple inter-cross breeding generated *Sptlc2* knockout mice (*Sptlc2*^lox/lox^.*Ins1*Cre^+/-^ referred as *Sptlc2*^ΔIns1^) and littermate controls (*Sptlc2*^lox/lox^.*Ins1*Cre^-/-^ referred as *Sptlc2*^CTL^). Experiments were performed in male.

### Mouse maintenance and diet

Animals were housed 2 to 5 per cage under controlled temperature (22°C) and light conditions (light/dark, 12 hr/12 hr) and were fed *ad libitum* with either standard laboratory chow diet (referred as CD) or high fat diet (referred as HFD) or ketogenic diet (referred as KD), starting at age 8 weeks and lasting for 6 weeks. At 14 weeks of age, animals were killed. Only male mice were studied.

### Metabolic phenotyping

Body weight, body mass composition and blood glucose levels (fed state) were measured weekly. Oral glucose tolerance test, OGTT or insulin tolerance test, ITT were performed after 6 h fasting, blood samples were collected from the tail (before gavage and at 15, 30, 60, 90 and 120 min). Plasma (for OGTT) was extracted at before and after glucose gavage (15 and 90 min) for insulin dosage by ELISA.

### Metabolic measurements

Serum leptin, MCP-1 and free fatty acids (FFAs) were determined by MILLIPLEX assays and colorimetric assay, respectively.

### In vitro insulin secretion from isolated islets

Mice were killed at 14 weeks of age and islets of Langerhans were isolated. Insulin secretion was measured during glucose-stimulated insulin secretion (GSIS) test, essentially as previously described in Amouyal and al. [24] with modifications as given in Supplemental Data in “Extended Methods”.

### Quantification of sphingolipids

Sphingolipid concentrations were determined in islets by LC-MS/MS methods. All details could be found in supplemental data in Extended Methods.

### Histological analysis

Insulin, glucagon and somatostatin staining were performed on pancreata extracted from deceased animal fed with CD of 14 weeks of age essentially as described [25] with modifications as given in Supplemental data in “Extended methods”.

### Measurement of intracellular free Ca^2+^, ATP/ADP and cell-cell connectivity

Functional multicellular Ca^2+^-imaging was performed as previously described [26], with modifications as given in Supplemental data in “Extended methods”.

Cytosolic ATP/ADP ratio changes in response to elevated glucose was measured after infection with the recombinant probe, Perceval [27].

### mRNA expression profiling (RNAseq)

RNA sequencing (RNA-Seq) data were processed as well as Gene Ontology (GO) enrichment analysis in islets from *Sptlc2*^ΔIns1^ and *Sptlc2*^CTL^, and detailed in Supplemental data in “Extended methods. The bulk RNA-seq data set is deposited in the Genome Expression Omnibus under accession numbers GSEXXXXX.

A previously published islet RNASeq data (NCBI GEO: GSE78183; [28]) from male mice fed a high fat (HF) or regular chow (RC) diet for 90 days was used to investigate the expression of *Sptlc1* and *Sptlc2* in response to metabolic stress. Normalised expression counts for the two genes were extracted from this dataset and boxplots were produced using Graph Pad Prism Version 10.4.1.

A previously published dataset (NCBI GEO: GSE164416; [29]) of human islets from partially pancreatectomised patients was used to measure expression differences of *Sptlc1* and *Sptlc2* between diabetic (T2D) and non-diabetic (ND) individuals.

### Quantitative RT-PCR

Total RNA was extracted from pancreatic islets using the miRNeasy micro kit (Qiagen). Reverse transcription was performed using Superscript IV and random hexamer (Promega). We performed a Quantitative PCR with SybrGreen and normalized each transcript by 18S expression to obtain relative quantification. Upon request, all primers can be provided.

### Statistical Analysis

Data are expressed as means ± SEM. Statistical significance was assessed using Student’s t - test and 1-way ANOVA or Mann and Whitney test and Kruskal-Wallis for non-parametric analysis (for non-normal distribution) to examine the effect of multiple variables. Two-way ANOVA was performed in OGTT and ITT analysis P-values <0.05 were considered significant. Statistical analysis was performed with GraphPad Prism (GraphPad Software version 10.4.1).

## RESULTS

### 1) Genetic deletion of *Sptlc2* in β-cells alters sphingolipid content

To induce *Sptlc2* deficiency exclusively in the pancreatic β-cell, we crossed *Sptlc2*^lox/lox^ mice to heterozygous animals expressing *Cre* under the control of the endogenous *Ins1* gene (Figure 1 A). This resulted in a substantial reduction of *Sptlc2* expression in islets from *Sptlc2*^ΔIns1^ mice (Figure 1 B). Lowered levels of *Sptlc2* Exon1 confirmed deletion (Figure 1 B). Ceramide content, and particularly that of long-chain (LC-Cer) Cer-16:0 and very long-chain ceramides (VLC-Cer) such as Cer-22:0, -24:0, -24:1, was reduced in islets of *Sptlc2*^ΔIns1^ mice under chow diet (CD) (Figure 1 C, suppl Figure 1 A). Under high fat diet (HFD), only VLC-Cer was reduced by *Sptlc2* deletion (Suppl Figure 1 B). The ratio of long chain to very long chain ceramide (LC/VLC-Cer) such as C16:0/C24:1; C18:0/C24:1, C20:0/C24:1 was increased in response to HFD (Suppl Figure 1 I and J). Moreover, lowering VLC-cer levels in islets of *Sptlc2*^ΔIns1^mice under each diet was associated with a further increase in this ratio (Suppl Figure 1 I et J). Derivatives of ceramide, such as sphingomyelin (SM) and hexosylceramide (Figure 1 C, D and suppl Figure 1 C, D, G, H) were also affected by *Sptlc2* deficiency in CD and HFD-treated mice to a similar extent to that of Cer (i.e. reduction of SM 16:0, SM 22:0, SM:24:0, SM 24:1). Interestingly, SM16:0 and HexoCer16:0 levels in HFD-treated *Sptlc2*^ΔIns1^ islets were also lowered dramatically (Supp Figure 1 D, H) suggesting that de novo ceramide synthesis is crucial to maintain complex sphingolipids levels in β-cell.

**Figure 1:**
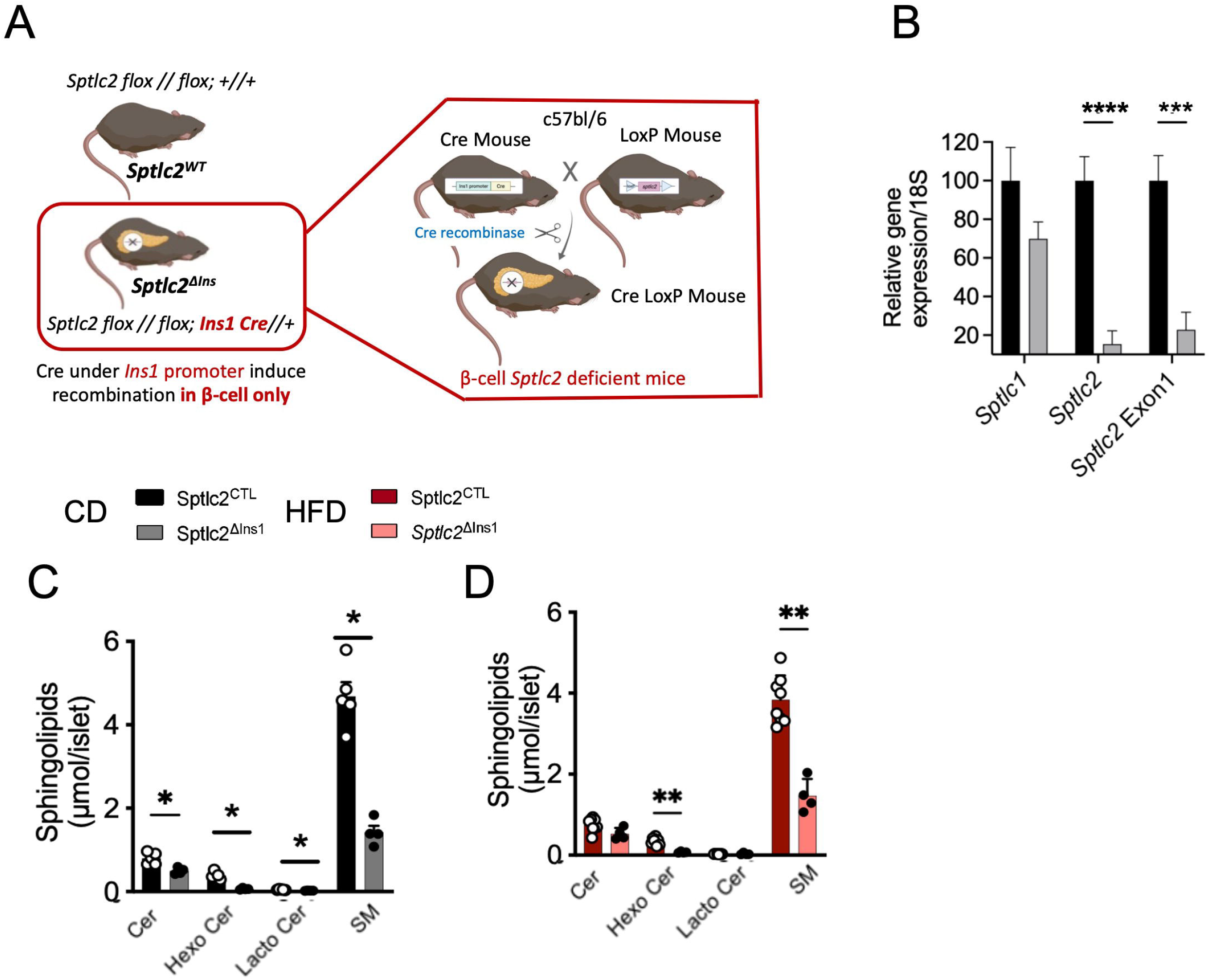
*Sptlc2* deficiency in β-cell modulate sphingolipids balance in pancreatic islet. Schematic representation of the recombination of the floxed alleles of *Sptlc2* using an Ins1-driven *Cre* recombinase (A). Gene expression of *Sptlc1, Sptlc2* and Exon1 of *Sptlc2* measured by qPCR in isolated islets of Sptlc2^CTL^ and Sptlc2^ΔIns1^ mice (B). Quantification of sphingolipids, ceramides and sphingomyelin in islets of regular chow (CD) and high fat (60% HFD) diet-fed mice (C-D). Analysis was performed in 14-week-old mice either on CD or HFD, starting at 8 weeks. Error bars represent SEM, n=4-8 mice per group, *p<0,05, **p<0,01, ***p<0,001 with Kruskal-Wallis or Mann and Whitney test.

### 2) Genetic deletion of *Sptlc2* impairs insulin secretion and glucose intolerance which are independent of diet

*Sptlc2* deficiency in β-cells did not affect body composition (Figure 2 A, B), though glucose homeostasis was drastically impaired in mutant animals on either diet. Firstly, fed glycemia was impaired from an early age (Figure 2 C, D). Furthermore, glucose tolerance tests (OGTT), performed at 12 weeks of age, revealed that *Sptlc2*^ΔIns1^ mice were markedly glucose intolerant compared to *Sptlc2*^CTL^ mice under both CD and HFD conditions (Figure 2, E, F, H, I). Plasma insulin increases during glucose gavage were eliminated at 15 min. in *Sptlc2*^ΔIns1^ mice on either diet (Figure 2 G and J). Importantly, insulin sensitivity was not affected by β-cell-selective *Sptlc2* deletion, and glucose excursions during insulin tolerance tests (ITT) were similar between genotypes for mice on CD (Fig 2 K, L) and HFD (Figure 2 M, N). Circulating levels of free fatty acids (Figure 2 O), leptin (Figure 2 P), and an inflammatory marker (MCP-1, Figure 2 Q) were not affected in *Sptlc2*^ΔIns1^ versus control mice, arguing against systemic inflammation or anomalies in systemic lipid metabolism.

**Figure 2:**
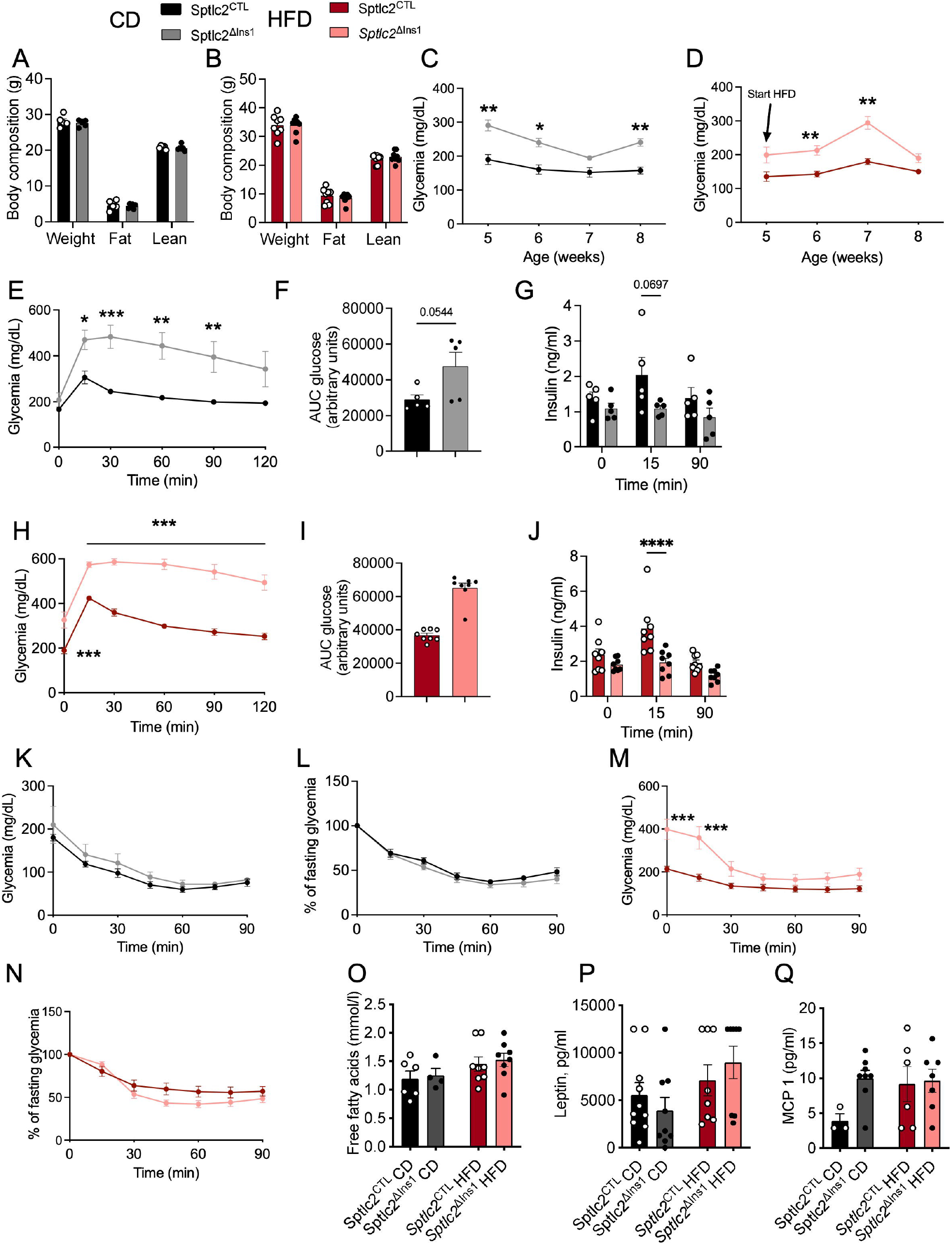
Impaired glucose induced insulin secretion in Sptlc2^ΔIns1^ mice regardless of the diet. Body composition (A, B). Morning fed glycemia of animals was followed from 5 to 8 weeks of age, under chow (C) or high fat diet initiated at 5 weeks of age (D). Oral glucose (2g/kg) tolerance test: glycaemia (E, F, H, I) and insulin secretion (G, J). Insulin (0.5 ui/kg) tolerance test for CD mice (K, L). The insulin dose was 0.75 iu/kg for HFD mice (M, N). Fasted metabolic analysis (free fatty acids, Leptin, MCP-1 (O-Q). Analysis was performed in 13 weeks old mice (except for the fed glycemia experiment) either on CD or HFD from 8 weeks of age. Error bars represent SEM, n=4-8 mice per group, *p<0,05, *p<0,01, ***p<0,001 with Kruskal-Wallis or Mann and Whitney test.

To remove a potential confounding effect of glucotoxicity on the regulation of carbohydrate metabolism of mutant mice, we fed mice with a ketogenic diet (KD) from weaning. At eight weeks of age, basal glycaemia and body weight were similar in *Sptlc2*^ΔIns1^ and *Sptlc2*^CTL^ mice (Supp. Fig 2 A-B). Although the ketogenic diet limited the glucotoxicity, glucose intolerance and substantial defects in insulin secretion persisted (Sup Fig 2 C-E). In addition, the sphingolipid content of islets from *Sptlc2*^ΔIns1^mice on KD showed the same characteristics as those of the same mice on CD and HFD, i.e. a significant reduction in the quantity of all Cer species and their derivatives in favor Cer (C18:0-Cer and C20:0-Cer) (Supp. Fig 1 F-H).

These results demonstrate that *Sptlc2*^ΔIns1^ mice are glucose intolerant with impaired insulin secretion which is independent of the metabolic state. Indeed, the phenotype is already severe in *Sptlc2*^ΔIns1^ mice under CD and is not reversed with the KD despite normal basal glycaemia.

Thus, the effects of *Sptl2* deletion are largely β-cell autonomous.

### 3) Sptlc2 deficiency in pancreatic β-cells leads to a drastic reduction in β-cell and islet number

Examined in mice on CD, total islet area was significantly reduced in *Sptlc2*^ΔIns1^ *versus Sptlc2*^CTL^ mice whereas total pancreas area was similar between the genotypes (Figure 3 A, B). β-cell deficiency of *Sptlc2* was associated with a particularly marked reduction in the number of small islets compared to that of control mice (Figure 3 C, D) and altered insulin-positive cell area (Figure 3 E-G). Glucagon-positive cell area was similar in both groups (Figure 3 H-J) whereas the proportion of somatostatin (SST) positive cells was significantly higher in islets of *Sptlc2*^ΔIns1^ *versus Sptlc2*^CTL^ mice (Figure 3 K-M).

**Figure 3:**
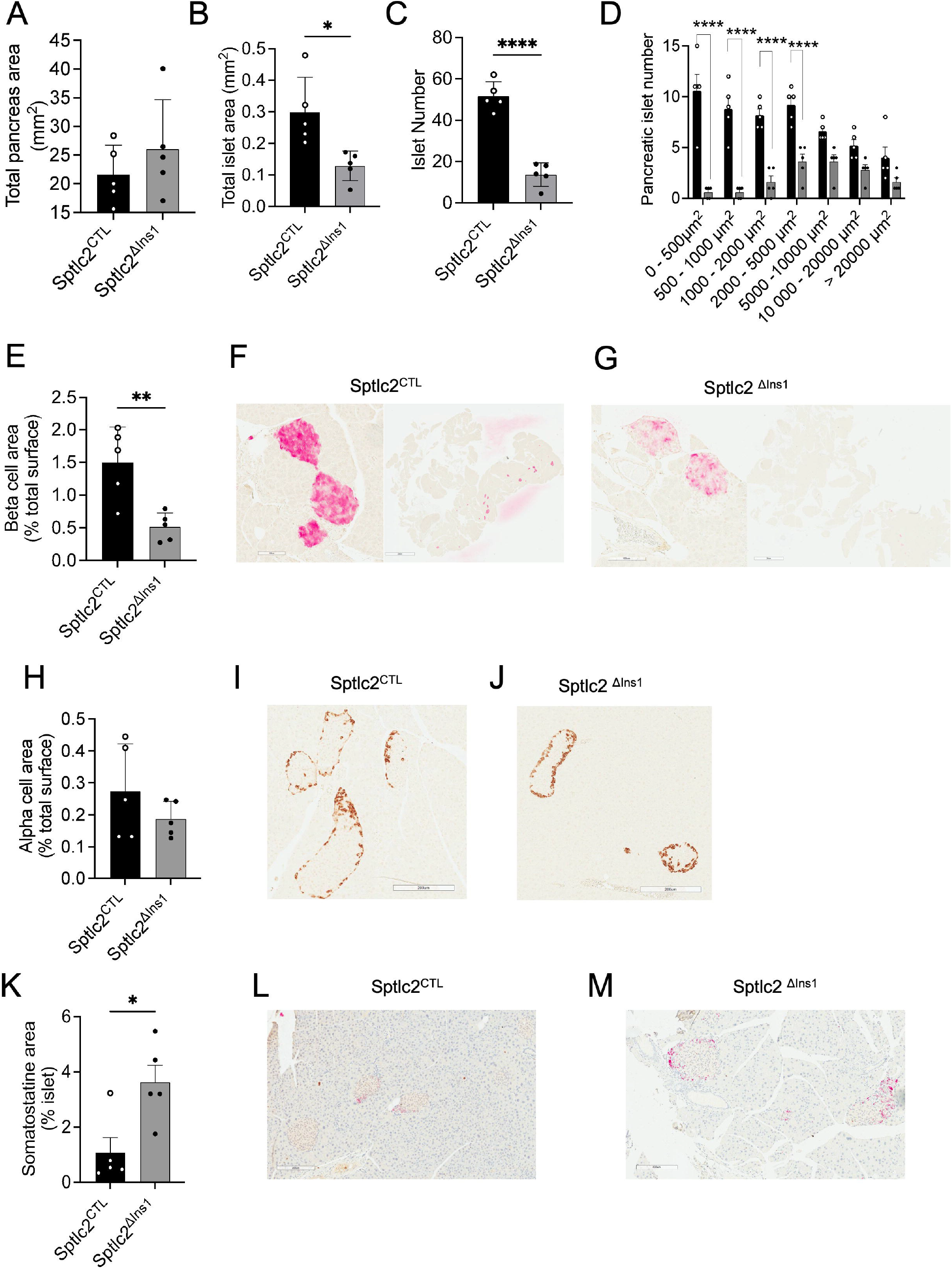
*Sptlc2* deletion in the β-cell induces a drastic reduction in the number of pancreatic islets. Pancreatic histology analysis for total area (A). Pancreatic histology analysis for insulin staining: evaluating total islets area (B), islets number (C-D) and β-cell area (E-G). Pancreatic histology analysis for glucagon (H-J) and somatostatin staining (K-M).

Somatostatin (*Sst*) gene expression was sharply increased in *Sptlc2*^ΔIns1^ mice versus controls under CD conditions, whereas *Gcg* expression was unchanged (RNAseq analysis, log2FC 1.22, padj 7.41E-04 and log2FC 0.97, padj 0.27 respectively).

Therefore, the marked impairment of glucose tolerance in *Sptlc2*^ΔIns1^ mice is likely to be the result of insulin deficiency reflecting a loss of β-cells and altered islet composition.

### 4) Pancreatic β-cells from *Sptlc2* deficient mice display preserved responses to glucose despite lowered insulin content and cellular stress

We next investigated islet function in *Sptlc2*^ΔIns1^ mice at 14 weeks of age. Examination of glucose-stimulated insulin secretion (GSIS) in pancreatic islets isolated from *Sptlc2*^ΔIns1^ or control mice (maintained under CD or HFD) revealed preserved hormone release in response to high glucose in each case (Figure 4 A, C). However, total insulin content was significantly reduced in *Sptlc2*^ΔIns1^ versus *Sptlc2*^CTL^ mice under both diets (Figure 4 B, D). In line with these results, imaging analyses revealed that islets from *Sptlc2*^ΔIns1^ and *Sptlc2*^CTL^ mice responded with similar oscillatory increases in intracellular free Ca^2+^concentration during stimulation with 17 mM glucose or 20 mM KCl (Figure 4 E, F). Similarly, increases in cytosolic ATP/ADP in response to high glucose were similar in the two groups, suggesting that mitochondrial oxidative activity is preserved in *Sptlc2*^ΔIns1^ mouse islets (Figure 4 G, H). In both groups, cell-to-cell coupling among β-cells, and the proportion of highly connected cells, increased similarly in response to an elevation in glucose from 3 to 17 mM. However, under KCl stimulation limited β-cell-β-cell de-coupling was observed in islets of *Sptlc2*^ΔIns1^ mice compared to those from *Sptlc2*^CTL^ animals (Figure 4 I-K).

**Figure 4:**
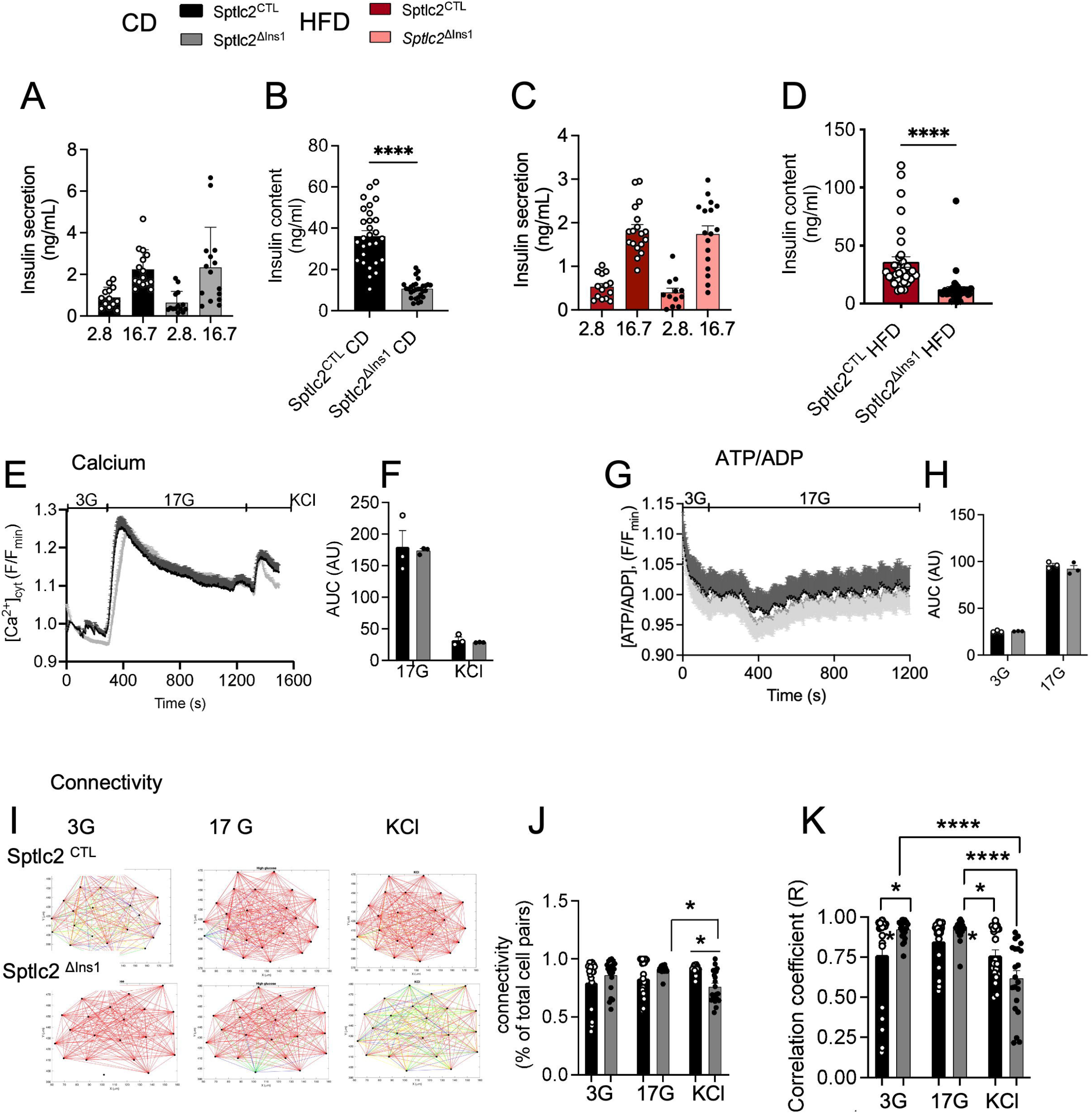
Preserved insulin secretory functionality despite β-cell-selective *Sptlc2* deficiency. Glucose-stimulated insulin secretion (GSIS) of isolated islets from 14-week-old mice under CD or HF was measured under low glucose (2.8mM) or high glucose (16.7mM) (A-C). Total insulin content (B-D). Measurement of intracellular free Ca2+ (E-F), ATP/ADP (G-H) and connectivity (I-K) assessed by imaging analysis under low glucose (3mM), high glucose (17mM) and KCl stimulation. Error bars represent SEM, n=3-5 mice per group, *p<0,05, **p<0,01, ***p<0,001 with Kruskal-Wallis or Mann and Whitney test.

Taken together, the above results demonstrate that the islets remaining in *Sptlc2*^ΔIns1^ mice under CD or HFD maintain normal functionality, including preserved glucose-induced insulin secretion.

In order to gain a better understanding of the β-cell defects observed in vivo, RNA sequencing (RNAseq) was performed. Comparison of the islet transcriptome of *Sptlc2*^ΔIns1^ and *Sptlc2*^CTL^ mice under CD and HFD identified 3125 and 8084 genes, respectively, that were differentially expressed in *Sptlc2*^ΔIns1^ compared to*Sptlc2*^CTL^ mice (adjusted p-value < 0.05) (Figure 5 A and B). Among the differentially expressed transcripts, cholinergic receptor muscarinic type 3 (Chrm3) mRNA was down-regulated in islets of *Sptlc2*^ΔIns1^ mice under CD, but not under HFD (log2FC -0.81, padj 7.8E-0.3 and log2FC -0.24, padj 0.35 respectively). These results suggest that de novo ceramide synthesis in β-cells could regulate insulin secretion in vivo by modulation of the parasympathetic action through the regulation of a well-known receptor that controls insulin secretion [30].

**Figure 5:**
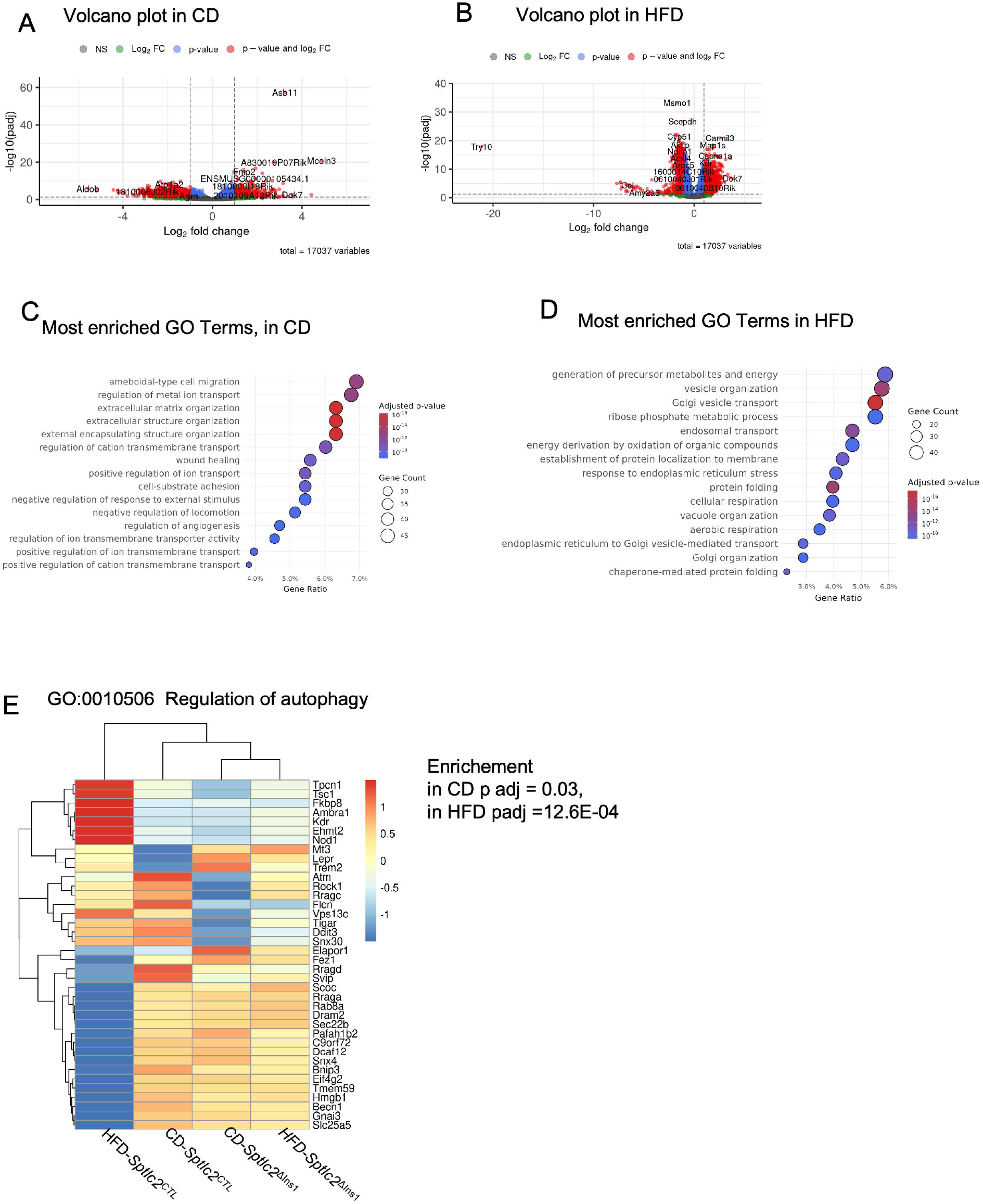
Alteration of membrane and autophagy function implied by transcriptomic analysis of islets from Sptlc2^ΔIns1^ mice. RNA seq analysis was performed using islets isolated from Sptlc2^ΔIns1^ or Sptlc2^CTL^ mice under chow (CD) and High fat (HFD) diet. Volcano plot of gene expression in CD (A) and HFD (B). Gene Ontology (GO) analysis of the most enriched terms in Sptlc2^ΔIns1^ compared to Sptlc2^CTL^ mice under CD (C) and HFD (D). Heatmap representing gene expression of enriched GO Terms: GO:0010506 Regulation of autophagy (E) n=3-4 mice per group, mice were aged of 14 weeks

To explore the latter findings in more depth, we performed a Gene Ontology (GO) Term enrichment analysis and identified 1491 and 589 biological processes significantly modified in *Sptlc2*^ΔIns1^ mice compared to controls on CD and HFD respectively (enrichment adjusted p-value ≤ 0.05). Among the 20 GO Terms most enriched in islets of CD *Sptlc2*^ΔIns1^ mice, seven referred to ion transport (Figure 5 C) whereas in islets of HFD *Sptlc2*^ΔIns1^ mice, five referred either to vesicle transport (Figure 5 D). In addition, autophagy-related biological functions (Figure 5E) were significantly enriched in *Sptlc2*^ΔIns1^ mice fed with CD and HFD diet. Also, essential protein recruited to phagosphore upon autophagy induction are significantly reduced in *Sptlc2*^ΔIns1^ mice fed with CD (*Ulk 2*: log2FC -0.48, padj 6.4E-0.3) and with HFD (*Ulk 1*: log2FC -0.68, padj 1.6E-0.4). These results suggests that membrane abnormalities impaired the autophagy capacity of pancreatic islet cells and probably β-cells.

In summary, despite preserved GSIS, islets from *Sptlc2*^ΔIns1^ mice show several defects: sharply lowered insulin content, membrane functional defects and impaired autophagy. Any of these defects could lead to β-cell loss observed by the pancreas histological analysis.

### 5) Changes in SPT2 expression in human and mouse islets in T2D

A transcriptomic analysis of islets isolated from 1292S, AJ, AKR, BALBC, C57Bl6 and DBA2J mice after 2, 10, 30 and 90 days of CD or HF diet [28] was re-interrogated. *Sptlc2* expression was reduced significantly after 30 days of HFD in BALBc and DBA2J mouse islets compared to islets of mice under CD (p=0.02 and p=9.8E-0.4 respectively) (Figure 6 D, F). Likewise, *Sptlc1* expression was reduced significantly after 10 days of HFD in BALBc and 30 days of HFD in DBA2J mice islets compared to islets of mice under CD (p=0.02 and p=2.4E-0.4 respectively) (Figure 6 J, L).

**Figure 6:**
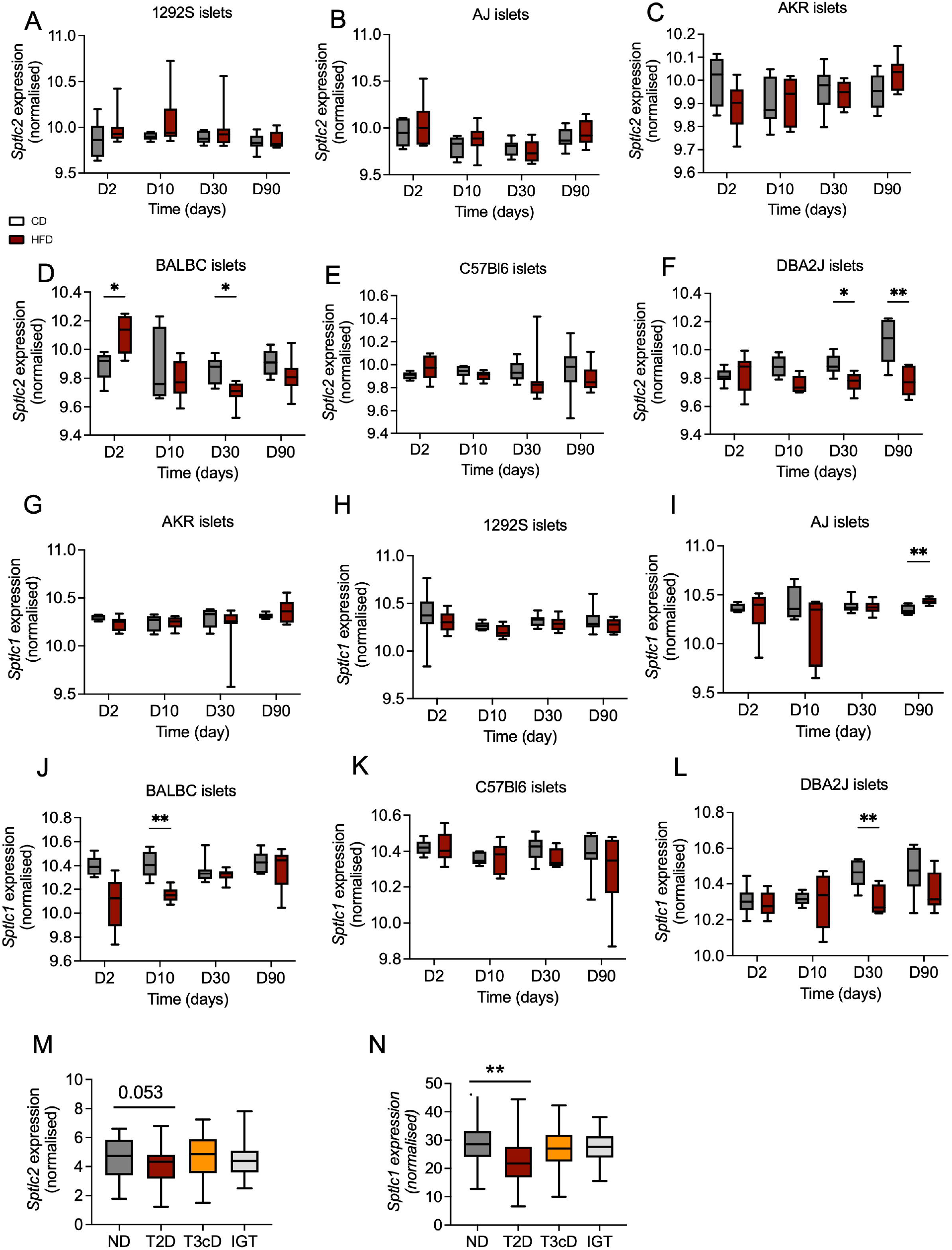
Reduced *Sptlc1* and *Sptlc2* expression induced by high fat diet in islets of BABLC and DBA2J mice. *Sptlc2* and *Sptlc1* expression in RNA seq analysis of isolated islets of 1292S (A, G), AJ (B, H), AKR (C, I), BALBC (D, J), C57Bl6 (E, K) and DBA2J (F, L) mice under chow diet (CD) and High fat diet (HFD) at different time point. n=6 male mice per group, mice aged of eight weeks. *Sptlc2* (M) and *Sptlc1* (N) expression in RNA seq analysis of isolated islets from pancreatectomized human non diabetic (ND) living with type 2 diabetes (T2D), diabetes secondary to pancreatic disease (T3cD) or glucose intolerant patient (IGT).

Finally, transcriptomic analysis was performed in human islets obtained from patients undergoing partial pancreatectomy [29]. *SPTLC2* expression tended to be lowered in islets from patients with T2D compared to non-diabetic controls (padj 0.053 and log2FC -0.27) and *SPTLC1* expression was significantly reduced (padj 3.5E-0.3 and log2FC -0.42) (Figure 6 M, N). In islets from patients with glucose intolerance or with diabetes secondary to pancreatic disease, *SPTLC1* and *SPTLC2* expression was not different from normogluco-tolerant patients (Figure 6 M, N).

## DISCUSSION

The present study demonstrates, firstly, the requirement for the *de novo* synthesis of ceramide for the maintenance of a normal β-cell mass and, secondly, the regulation of both *Sptlc1* and *2* by nutrients. Further, we demonstrate that changes in either or both SPT isoforms may be involved in the pathogenesis of T2D.

Ceramide accumulation is a likely contributor to tissular insulin resistance in obesity and T2D [4]. Recent evidence has suggested a beneficial role for ceramides species in some metabolically-sensitive tissues [31] and β-cell for normal glucose homeostasis [18]. In agreement, in our present study, we show that blockade of *de novo* ceramide synthesis through deletion of *Sptlc2* is deleterious towards β-cells. Thus, deletion of *Sptlc2* using an endogenous *Ins1* gene-drive *Cre* [23] results in the loss of ∼80 % of β-cells.

Our findings emphasize the critical importance of preserved sphingolipid including long chain ceramide and its derivatives (SM and glycosphingolipid) in the β-cell, presumably reflecting essential roles of the latter in maintaining membrane integrity and intracellular signaling [15]. Recent studies have suggested that specific ceramide species produced through CerS2 control proinsulin processing [18]. In accordance, we found that *Sptlc2* deletion also induced a drastic decrease of insulin content in islets. Griess et al proposed that processing of proinsulin is reduced in the absence of sphingolipid produced by CerS2 through a post-translational mechanism [18]. Other studies have evidenced a central role of ceramide derivatives such as sulfatides in insulin processing [32]. Immuno-electron microscopy also revealed an intracellular expression of sulfatide in the secretory granule [33]. Our lipidomic screen showed that *Sptlc2* deletion drastically decreased HexosylCer, which are the precursors of sulfatides, supporting the idea that *de novo* ceramide synthesis is crucial to maintain the secretory granule membrane in β-cells. Furthermore, SM levels in *Sptlc2*^ΔIns1^mice, which are an essential component of membranes, are also drastically decreased. In line with this, expression of *Smpd5* (encoding the major sphingomyelinase in the pancreas) was up-regulated in islets of *Sptlc2*^ΔIns1^mice under CD (RNAseq analysis, log2FC 1.96, padj 0.035). Expression of *Smpd1* and *Smpdl3a* were also up-regulated in islets of Sptlc2^ΔIns1^mice under HFD (RNAseq analysis, log2FC 0.484.12, padj 3.14E-04 and log2FC 0.65, padj 7.5E-05 respectively). Thus, in animals fed either diet, inhibition of *de novo* ceramide synthesis in β-cells leads to compensatory strategy to maintain Cer content involving the sphingomyelinase pathway. Nevertheless, these are insufficient to prevent the observed lowering of Cer and SM.

Deletion of sphingomyelin synthase 1 (SMS1), which catalyzes the production of SM, has been shown to impair insulin secretion [34] due to mitochondrial dysfunction. However, despite a drastic decrease of SM levels in *Sptlc2*^ΔIns1^ islets, *ex vivo* insulin secretion by *Sptlc2*^ΔIns1^ islets in response to glucose was not altered, suggesting a potential compensatory mechanism.

In addition to the regulation of insulin processing/secretory granule formation, deletion of *Sptlc2* and the associated decrease of ceramide and its derivatives appears to be a key regulator of β-cell mass. In line with our results, Castell et al. recently showed that oleate infusion, which increases very long-chain unsaturated ceramides (Cer 24:1), promotes β-cell proliferation via activation of de novo ceramides synthesis [35]. Ceramides and their derivatives have not yet been associated with β-cell differentiation. However, Hua and colleagues have recently shown that ceramide metabolism is also crucial for functional maturation of human pluripotent stem cell-derived β-cells [36]. Moreover, it has been proposed that synthesis of glycosphingolipid structures is essential for embryonic development and for the differentiation of some tissues such as neuronal tissue [37]. Future experiments will be required to determine the effect of ceramide and its derivatives on β-cell differentiation/proliferation.

In order to gain a fuller insight into the role of *de novo* ceramide synthesis for β-cell development, we performed a transcriptomic analysis. Although the remaining β-cells remain normally glucose-responsive *ex vivo*, substantial changes were evident at the transcriptomic level and these may influence insulin secretion or survival *in vivo*. Deletion of *Sptlc2* in β-cells was not associated with an alteration of genes involved in β-cell differentiation, identity and function (Suppl Figure 3). These results support the idea that decrease of ceramides and its derivatives will probably alter membrane functions or dynamics that impact β-cell fate. Indeed, membrane dynamics are central during development [38]. Among the most highly altered pathways in *Sptlc2*^ΔIns1^ islets correspond to the regulation of ion transport and vesicular transport, which both rely on the integrity of cellular membranes. Looking more carefully at GO pathways related directly to membrane properties, islets from *Sptlc2*^ΔIns1^ mice exhibit an alteration in the regulation of autophagy. Inside the GO “regulation of autophagy”, a cluster of genes was down-regulated in response to *Sptlc2* deletion. Previous studies have shown that loss of autophagy decreased β-cell mass and pancreatic insulin content due to an increased apoptosis and decreased proliferation of β-cells [39]. Moreover, ceramides are critical regulators of autophagy [40] supporting the idea that inhibition of de novo ceramide synthesis will decrease β-cell mass by impairing autophagy. In HFD conditions, data are contrasting since a cluster of genes “regulation of autophagy” are upregulated whereas another are down-regulated. However, deletion of *Sptlc2* completely abrogated these regulations, supporting a role of de novo ceramide synthesis.

Finally, looking individually at differentially expressed genes in islets from *Sptlc2*^ΔIns1^ mice, the *Chrm3* encoding the cholinergic receptor muscarinic type 3, was significantly decreased. In addition to its well-known role in the control of insulin secretion by the parasympathetic axis, it has been proposed that this axis could control β cell mass establishment in the postnatal period [30]. It is therefore conceivable that regulation of *Chrm3* by *de novo* ceramide synthesis is an important regulator of β-cell development.

Suggesting that the present data in mice may be relevant for human T2D, variants in the *SPTLC1* locus, encoding the non-redundant SPT1 subunit of the complex, are moderately (HuGE score 8.4, 3.0 respectively) [41, 42] associated with HOMA-B and with fasting insulin, with weaker associations in this region for T2D. Levels of SPTLC1 protein also tend (adjusted p-value, 0.118) to be decreased in islets from subjects with diabetes versus controls [43]. A lowering in *SPTLC2* mRNA was reported in islets from human subjects with T2D [44], and with newly diagnosed type 1 diabetes (T1D) [45]. Our own data in human subjects are in line with those findings. However, more recent studies [43, 46] failed to replicate these results, and alterations in *SPTLC2* mRNA were not apparent. Nevertheless, we show that islet *Sptlc2* and *Sptlc1* mRNA levels are lowered by high fat diet treatment in two mouse lines (BALBc and DBA2J) demonstrating nutrient-responsive regulation of this gene in preclinical models. Studies of β-cell selective *Sptlc1* null mice, once available, are thus likely to be informative.

An interesting finding of the present studies was the 2-3-fold increase in islet somatostatin gene expression, and in the total number of delta-cells in the pancreas of *Sptlc2*^ΔIns1^ mice. These changes are likely, at least in large part, to reflect a lowering in β-cell number, as previously reported in models of T1D including nonobese diabetic (NOD) or streptozoticin-treated mice [47] (though others have reported no changes in delta cell number in NOD mice; [48]). In addition, it is possible that functional changes in the remaining β-cells (e.g. in autophagy regulation) influence the synthesis and release of undefined factors from these cells post-natally or in the adult. Of note, the latter changes in delta-cell number in T1D models were not ascribed to hyperglycemia in the study by Plesner and colleagues, and were unaltered by the normalisation of blood glucose though islet transplantation [47]. The observed increase in delta:beta cell number may thus contribute to altered in vivo insulin secretion and may also influence glucagon secretion, as reported recently [48].

In contrast to the above T1D models, the loss of β-cells after *Sptl2* deletion appears to be cell autonomous, thus resembling similar changes after the deletion of other genes critical for β-cell survival [49]. Correspondingly we observed no evidence for insulitis nor of transcriptomic alterations indicative of immune attack (e.g. cytokine response gene induction) [50]. Thus, *Sptl2*^ΔIns1^ mice provide a model with features of both T1D and T2D.

An important conclusion of our results is that global blockade or inhibition of SPT2, achieved with pharmacological approaches which seek to rescue insulin sensitivity in T2D [9], is likely to be deleterious at the level of the pancreatic β-cell, and thus to suppress insulin production. Such treatment strategies should therefore be approached with caution, until such time as they can be restricted to selected cell types (e.g. liver or skeletal muscle) through theragnostic strategies.

## ACKNOWLEDGMENTS

We are grateful to Mélanie Guévremont and the CRCHUM Cellular Physiology core facility for islet cell mass analyses.

## CONTRIBUTION

CA, SPC, JD, IR, NK, MD, FM, JL, EG, MI, MC performed the experiments, CA, HLS and GAR wrote the manuscript, CCG, HLS, GAR, CM reviewed the manuscript.

