## Supplementary figures and images for "Serine palmitoyltransferase-mediated de novo sphingolipid biosynthesis is required for normal insulin production and glucose tolerance"

### Supp 1

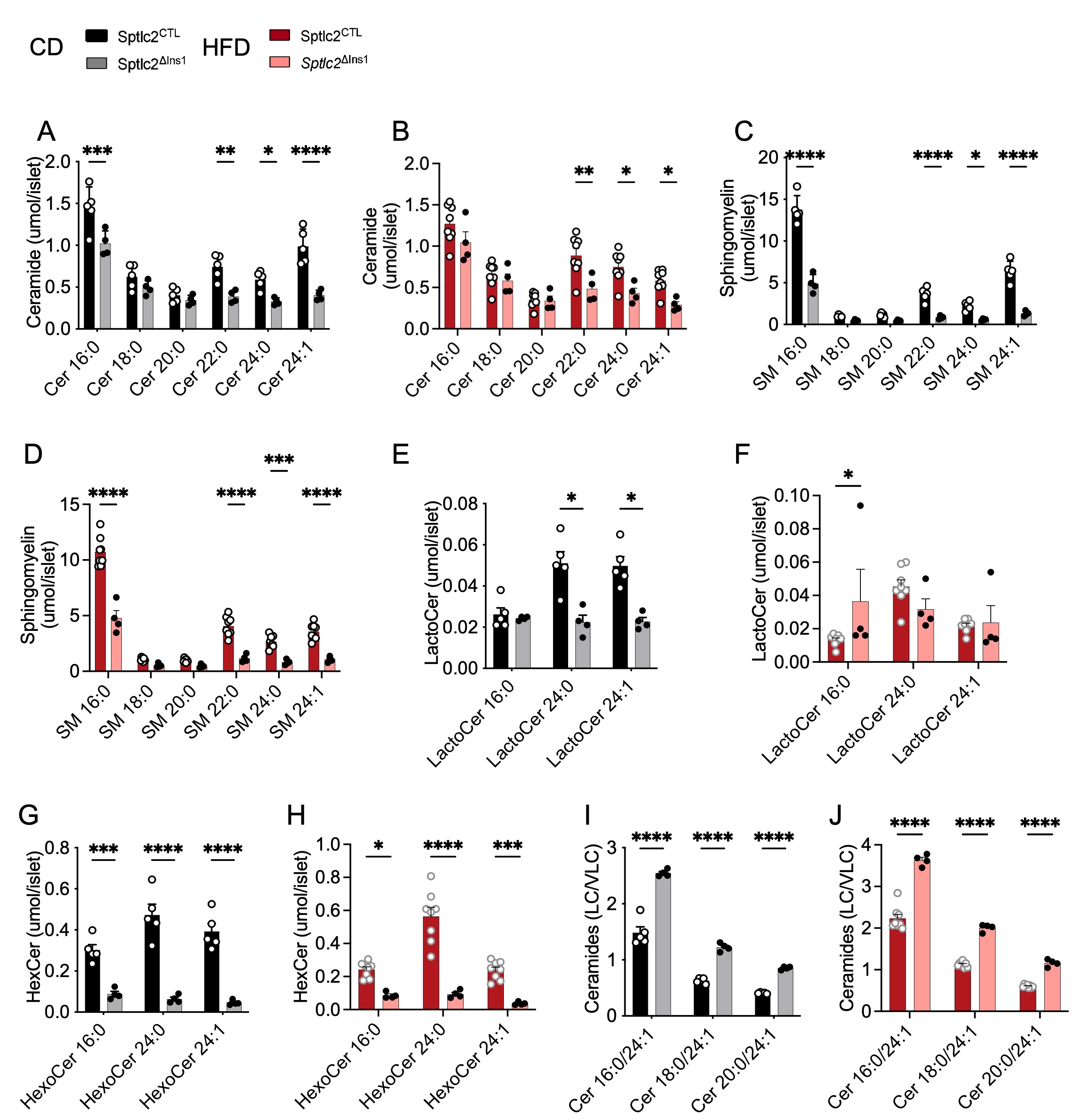

### Supp 2

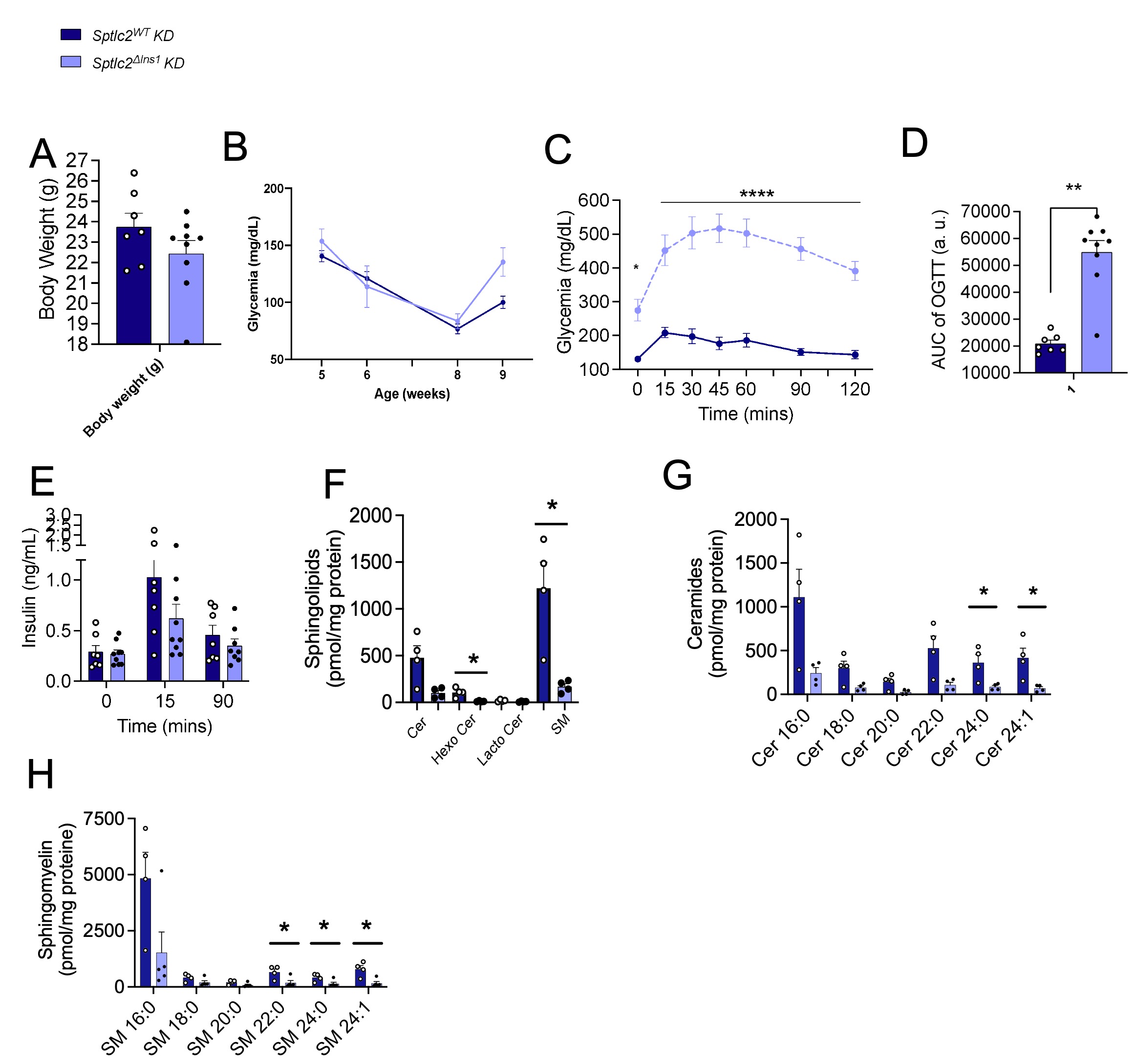

### Supp 3

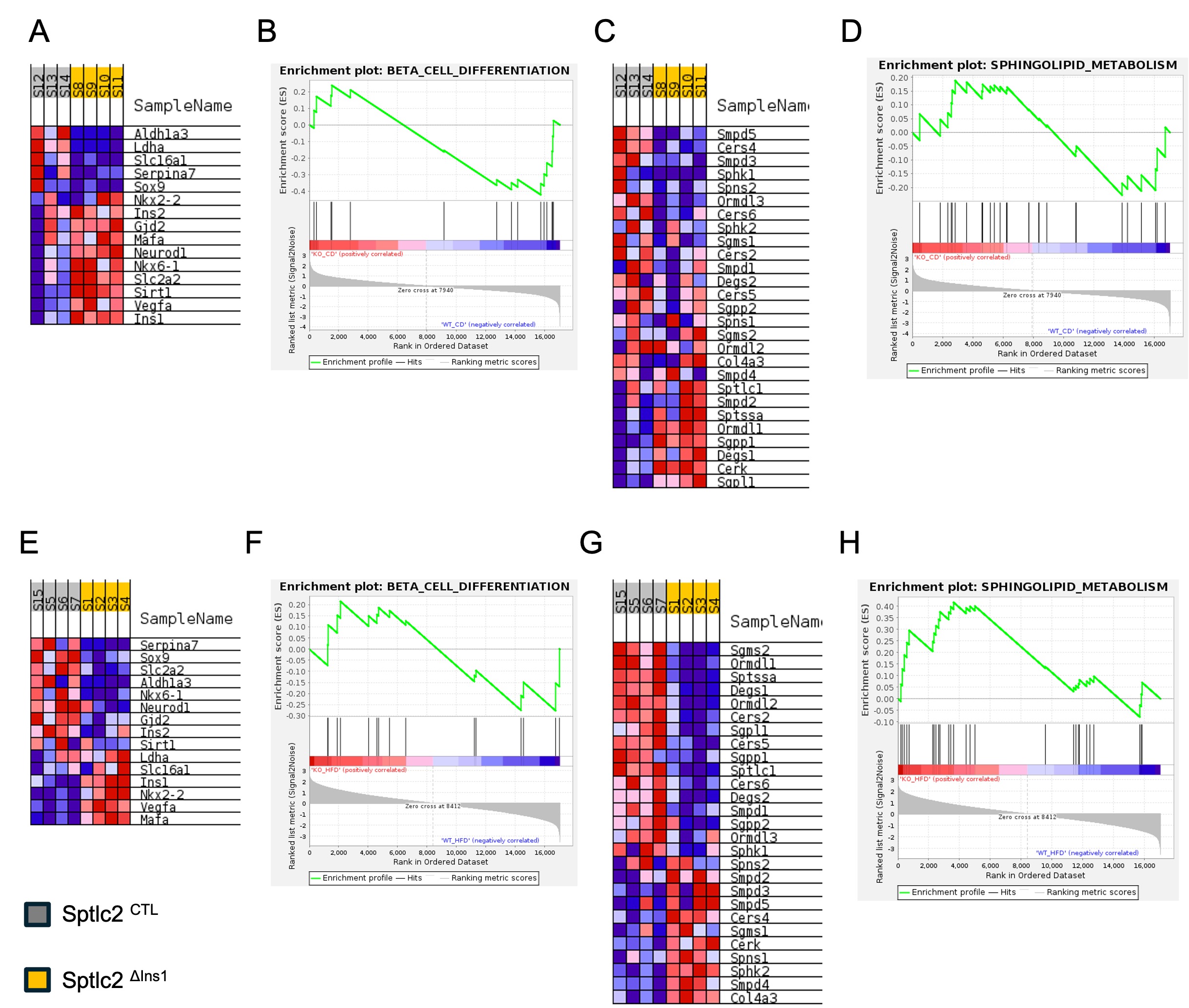
